# miR-18a-5p upregulates Orai1 expression to promote vascular smooth muscle cell proliferation and neointimal hyperplasia after injury

**DOI:** 10.64898/2026.01.20.700592

**Authors:** Marta Martín-Bórnez, Beltzane Dominguez-Liste, Javier Ávila-Medina, Eva Sanchez de Rojas-de Pedro, Isabel Galeano-Otero, José Sanchez-Collado, Antonio Ordoñez, Juan Antonio Rosado, Abdelkrim Hmadcha, Debora Falcon, Tarik Smani

## Abstract

**Rationale:** Neointimal hyperplasia, a key contributor to restenosis, is driven by the abnormal proliferation and migration of vascular smooth muscle cells (VSMC), although the underlying molecular mechanisms remain incompletely understood. This study aimed to characterize the structural, transcriptomic, and post-transcriptional changes driving neointima formation, with a focus on store-operated calcium entry (SOCE) pathways and microRNA (miRNA) regulation.

**Methods:** A rat carotid angioplasty model was employed to assess neointimal development at 1, 2, and 3 weeks post-injury. VSMC isolated from rat coronary arteries and A7r5 VSMC line were used to assess intracellular Ca^2+^ dynamics, and expression of gene and protein.

**Results:** Progressive neointimal thickening and impaired contractility were observed after carotid artery injury, accompanied by significant VSMC proliferation. Transcriptomic profiling revealed differentially expressed genes (DEGs) at 1 and 3 weeks, with enrichment in pathways related to cell cycle, migration, and Ca^2+^ signaling. Among Ca^2+^ -regulatory genes, Orai1 and SARAF were upregulated in the neointima and shown to colocalize and interact post-injury. Functional studies in VSMC demonstrated that Orai1, but not SARAF, is involved in insulin-like growth factor 1 (IGF-1)-induced proliferation and SOCE activation. Moreover, miRNA profiling identified miR-18a-5p, from the miR-17-92 cluster, as the most upregulated miRNA early post-injury. miR-18a-5p unusually enhanced Orai1 promoter activity and protein expression, leading to increased SOCE in VSMC. In human coronary arteries from ischemic hearts Orai1 was upregulated, suggesting the potential translational relevance of Orai1 in vascular pathology.

**Conclusions:** Our findings identify miR-18a-5p as a novel positive regulator of Orai1 and SOCE activity in VSMC. The results uncover a miRNA/SOCE regulatory circuit that orchestrates Ca^2+^ -dependent VSMC signaling during vascular remodeling and may serve as a potential therapeutic target pathway in occlusive vascular disease.

## INTRODUCTION

Percutaneous coronary intervention (PCI) and stent implantation are widely used surgical interventions for treating atherosclerotic vascular diseases, particularly coronary artery disease. However, high restenosis rates have been reported in patients who have undergone successful coronary angioplasty, due to vascular remodeling and neointimal hyperplasia (*1*, *2*). In fact, despite significant advances made with improved drug-eluting stents, releasing antiproliferative drugs, neointimal hyperplasia remains a major challenge for sustained therapeutic success (*3*). Therefore, understanding the mechanisms underlying balloon injury-induced neointimal hyperplasia is essential for developing new approaches to mitigate this pathological process.

Vascular smooth muscle cells (VSMC) proliferation and migration play pivotal roles in the pathogenesis of restenosis, which is associated with phenotypic switching of VSMC from a contractile to a synthetic phenotype induced by wide range of pathological stimuli. This switching stimulates the proliferation and migration of VSMC from the tunica media to the intima of the vessel wall, leading to uncontrolled neointimal hyperplasia. While the precise underlying mechanisms remain unclear, dysregulated intracellular Ca^2+^ signaling plays a crucial role in VSMC phenotypic switching. Among the relevant Ca^2+^ pathways, Store-Operated Calcium Entry (SOCE) is a crucial mechanism for Ca^2+^ influx in VSMC proliferation and migration within the neointima. The expression of key players of SOCE as STIM1, the Ca^2+^ sensor, and Orai1, the ion channel subunit, is upregulated in proliferative VSMC and in the neointima after vessel lesion (*4*, *5*). In contrast, SOCE-associated regulatory factor (SARAF), which modulates STIM1 activation (*6*, *7*), appears to be downregulated in balloon injury-induced neointima, correlating with reduced VSMC proliferation and migration (*8*). However, the molecular mechanisms involved in the expression of these proteins during neointima formation is not fully understood.

Mechanisms driving the upregulation of specific genes following vascular injury are complex and can be multifactorial. While transcriptional regulation has been extensively studied, emerging evidence suggests that post-transcriptional mechanisms, particularly microRNAs (miRNAs), play a pivotal role in fine-tuning gene expression in response to pathological stimuli. miRNAs primarily function by repressing gene expression post-transcriptionally through mRNA silencing or cleavage (*15*). But, under specific cellular contexts and conditions, miRNAs can also upregulate gene expression through distinct transcript and protein interactions (*16*). miRNAs act as significant modulators in regulating VSMC pathologies, including phenotypic switch, proliferation, and migration (*9*, *10*). However, the specific miRNAs responsible for modulating key genes involved in regulating Ca^2+^ signaling or VSMC proliferation, within vascular remodeling context, remains poorly defined. Only few studies analyzed the regulation of SOCE components by miRNAs in VSMC. For instance, miR-424/322 has been shown to indirectly regulates STIM1 during neointima formation in a rat angioplasty model (*11*). Meanwhile, Orai1 regulation by miRNAs has been investigated in cell lines, such as HeLa (*12*), T cells (*13*), or in mesangial cells (*14*), but it has not yet been explored in VSMC.

Given the critical role of SOCE and its potential for post-transcriptional control, we hypothesized that specific miRNAs dysregulated after vascular injury directly modulate SOCE components to drive VSMC proliferation. Therefore, we analyzed miRNAs dysregulation during neointima formation using a carotid artery balloon-injury rat model, and we investigated whether SOCE proteins can be modulated by miRNAs in VSMC in this context.

## METHODS

### Study setting and approval

All experiments involving animals adhered to the Royal Decree 53/2013, in compliance with Directive 2010/63/EU of the European Parliament and were approved by the Ethics Committee on human Research of the University Hospital of Virgen del Rocio of Seville (Permission number: CEI 2013PI/096) and the Animal Research Committee of the University of Seville (CE 27-05-256). Written informed consent was obtained from all patients.

### Human artery dissection

Human epicardial coronary arteries (left anterior descending) were dissected from the hearts of patients with end-stage heart failure (NYHA stage III–IV) due to ischemic dilated cardiomyopathy who were undergoing orthotopic heart transplantation, as described previously (*17*). As healthy controls, we used human carotid arteries isolated from donors after removal of other organs destined for transplantation, such as the heart.

### Rat carotid angioplasty-induced injury model

Adult male Wistar rats weighing 350–400 g were used and kept in optimal housing conditions, at 23-27°C with a 12 light/dark cycle, and were given free access to food and water before and after surgery. Rat carotid artery injury, sacrifice, and tissue collection were performed as previously described (*18*). Briefly, they were anesthetized with a mixture of 2% O_2_/sevoflurane and then with an intraperitoneal (i.p.) mixture of 50 mg/kg ketamine hydrochloride plus 20 mg/kg xylazine hydrochloride. During the surgical procedure, sedation was monitored by periodic observation of respiration and response to pain stimuli. The procedure was performed in the right common carotid artery (CCA). Given that vascular injury induces inflammatory responses (*19*), the left carotid artery from the injured animal model was used as internal control. The right carotid artery was isolated, and blood flow was interrupted using microclamps placed on both the internal and common carotid arteries. Then, a catheter with a Fogarty 2F balloon (Edwards Lifesciences, Irvine, CA 92614US) was inserted into the CCA through an incision made in the external carotid artery, and the balloon was inflated 3 times at a pressure of 1.5 atm to induce vascular endothelial injury. Finally, the external carotid artery was ligated, and blood flow was restored by releasing the microvascular clamps on the internal and common carotid arteries. After recovery from surgery and anesthesia, rats were subcutaneously injected with meloxicam (1 mg/kg) for pain management. Rats were sacrificed 24 h, 1 week (1W), 2 weeks (2W) or three weeks (3W) post-surgery via intraperitoneal (i.p.) administration of a lethal dose of sodium thiopental (200 mg/kg). Tissues were collected and harvested for hematoxylin-eosin, and immunofluorescence staining of cross-sections or protein, RNA, and microRNA analysis.

### In vivo analysis with ultrasound imaging

The Carotid Intima-Media Thickness (CIMT) and carotid artery shortening were measured using B-mode imaging on the Vevo^TM^ 2100 ultrasound system with a linear MS250 transducer with a frequency range of 13-24 MHz (VisualSonics, Toronto, ON, Canada). CIMT and artery shortening were analyzed pre- and postoperatively in anesthetized rats using 2% sevoflurane. The respiratory rate, heart rate, and core body temperature were monitored. A water heater and a heat lamp were used to maintain the temperature at approximately 37°C. As shown in video 1 and 2 in the supplemental data, near and far wall CIMT is clearly visible, and end-diastolic longitudinal images were acquired for at least three consecutive beats and were measured manually in a digitized scale.

### Morphometric analyses of carotid arteries

Rats were sacrificed 1, 2, or 3 W after surgery. Control and injured carotid arteries were extracted, cleaned of blood in 1X Phosphate Buffered Saline (PBS 1X) and fixed in formalin solution (formalin neutral buffered, 10% overnight. To evaluate vascular wall thickening, fixed samples were embedded in paraffin. Carotid sections were prepared in a motorized microtome (Leica CM1950, Wetzlar, Germany) at 10 µm and hematoxylin-eosin staining was performed according to standard protocols. Images were captured using a microscope BX51 Olympus (Olympus Soft Imaging Solutions, Japan) equipped with a DP70 digital camera. Objectives of 10X, 20X and 40X magnifications were used, and images were analyzed using CellP software (Olympus Soft Imaging Solutions, Japan) and ImageJ software (NIH, Bethesda, MD, US). The neointima, media, and internal elastic area were measured to calculate the neointima/media ratio, as well as the combined area of the intima and media layers called CIMA (Carotid Intima-Media Area).

### Immunofluorescence analyses

For immunofluorescence staining, fixed carotid arteries sections were placed in 15% sucrose followed by 30% sucrose in PBS 1X overnight. They were embedded in OCT (Optimum Cutting Temperature) compound Sakura Finetek, Torrance, CA, US) and stored at −80°C. Afterwards, carotid sections were permeabilized and blocked with 10% goat serum and 3% bovine serum albumin in phosphate-buffered saline (PBS1X) with 0,1% Triton for 1 h. After blocking, the slides were incubated with primary antibodies: rabbit anti-CD31 (1:400, Novus Biologicals, US), mouse anti-αSMA (1:400, Sigma-Aldrich, San Luis, MI, US), rabbit anti-SARAF (1:200, Abcam, Cambridge, UK) or mouse anti-Orai1 (1:200, Novus Biologicals, Centennial, CO, US) at 4°C overnight. Sections were then incubated with species-specific secondary antibodies conjugated to Alexa Fluor 488 goat anti-mouse-IgG and Alexa Fluor 594 goat anti-rabbit-IgG (Cell Signaling, Danvers, MA, US) for 4 h. Tissues were washed with PBS, incubated with DAPI (1:1000, Sigma-Aldrich, San Luis, MI, US) for 5 min to visualize nuclei, and mounted in DAKO fluorescence mounting medium (Agilent Technologies, Santa Clara, CA, US). Signal fluorescence was visualized and analyzed at the same z-plane in a Leica TCS SP2 using a 10X and 20X objective and the images were analyzed using ImageJ software (NIH, Bethesda, MD, US). Fluorescence intensity was quantified using LAS AF (Leica Application Suite; Leica Microsystems, Wetzlar, Germany).

For the spatial colocalization analysis of SARAF and Orai1 after vascular injury, proximity ligation assay (PLA) was performed using a Duolink *in situ* PLA detection kit (Sigma-Aldrich, St Louis, MO, US), as described previously. Briefly, sections from control and injured arteries were incubated with primary antibodies: anti-Saraf (1:200 rabbit) and anti-Orai1 (1:200 mouse) in antibody diluent overnight at 4°C. Proteins were labeled with Duolink PLA anti-rabbit PLUS and anti-mouse MINUS probes for 1 h at 37°C. Hybridization and amplification were observed when the proteins were in close proximity (<40 nm), detecting red fluorescent signals. Images were obtained at 10, 20 and 63X using maximum intensity projections of all z-sections (0.4 μm).

To analyze cell proliferation in carotid sections we used mouse anti-Ki67 (1:50, Thermo Fisher Scientific, Waltham, MA, US). Three randomly selected fields from three different rats from each group (1W, 2W, and 3W) were quantified as the percentage of Ki67-positive cells relative to the total number of cells stained with DAPI. Ki67-positive cells were counted using the FIJI/ImageJ Cell Counter plugin, whereas nuclei (DAPI-positive cells) were automatically counted using ImageJ.

### Culture of rat coronary vascular smooth muscle cells and A7r5

Coronary smooth muscle cells were isolated from the second branch of left anterior descending (LAD) coronary artery. The primary culture of VSMC was prepared following the same protocol as described previously (*20*). Briefly, the rat heart was quickly removed and placed in cold physiological solution (137 mM NaCl, 5.4 mM KCl, 0.2 mM CaCl_2_, 4.17 mM NaHCO_3_, 2 mM MgCl_2_, 0.44 mM KH_2_PO_4_, 0.42 mM NaH_2_PO_4_, 10 mM HEPES, 11.11 mM Glucose and 0.05 mM EGTA; pH 7.4) for coronary dissection. The LAD arteries were then incubated with 1.5 mg/ml elastase (4 units/mg) and 4 mg/ml collagenase type I (125 units/mg) (Sigma-Aldrich, San Luis, MI, US) in Dulbeccós Modified Eaglés Medium (DMEM) for 1 h at 37°C. Cells were mechanically dispersed and plated in DMEM supplemented with 10% fetal bovine serum (FBS, Thermo Fisher Scientific, Waltham, MA, US) and 100 U/ml penicillin and streptomycin, in a humidified atmosphere containing 5% CO_2_ and 95% air at 37°C. VSMC were easily distinguished by their size and typical elongated shape. To confirm the purity of the VSMC preparation and exclude any significant presence of fibroblasts or endothelial cells, the cells were stained with mouse anti-α-SMA antibody (Sigma-Aldrich, San Luis, MI, US). In the case of rat aortic A7r5 VSMC (CRL-1444, ATCC, Manassas, VA, US), they were expanded in basal media (DMEM supplemented with 10% FBS) and 100 U/ml penicillin and streptomycin. For cytosolic Ca^2+^ measurements and immunostaining, cells were plated on coverslips.

### Determination of the intracellular free intracellular calcium concentration ([Ca^2+^]_i_)

Cells were loaded with 2.5 μM Fura-2AM and imaged using an inverted microscope Leica (Wetzlar, Germany) equipped with a 20X fluor objective (0.75 N.A.), a monochromator (Polychrome V, Till Photonics, Munich, Germany), and a light-sensitive CCD camera (Cooke PixelFly, Applied Scientific Instrumentation, Eugene, OR), controlled by HP software (Hamamatsu Photonics, Shizuoka, Japan), as previously described (*21*). Experiments were performed under continuous perfusion in a Ca^2+^-free solution (120 mM NaCl, 4.7 mM KCl, MgCl_2_, 0.2 mM EGTA, 10 mM HEPES; pH 7.4), and the Ca^2+^ influx was determined from changes in Fura-2 fluorescence after re-addition of Ca^2+^ (70 mM NaCl, 2 mM CaCl_2_, 70 mM KCl, 1 mM MgCl_2_, 10 mM HEPES, 10 mM Glucose; pH 7.4). IGF-1 (10 ng/ml) or thapsigargin (TG) (2 μM) -induced Ca^2+^ influx was evaluated under different conditions, and represented as the ratio of Fura-2 fluorescence at an emission wavelength of 510 nm due to excitation at 340 and 380 nm (ratio = *F*_340_/*F*_380_). Δratio was calculated as the difference between the peak ratio after extracellular Ca^2+^ addition and its level immediately before the addition. In some experiments, GSK 7975A (GSK,10 μM) (Aobious, Gloucester, MA, US) was applied at the end of the experiment to inhibit SOCE.

### Cell transfection

VSMC were transfected with pre-designed small interference RNA (siRNA) at 70% confluence according to the manufacturer’s protocol using HiPerFect Transfection Reagent (Qiagen, Hilden, Germany). siOrai1, siSARAF or siControl (scrambled RNA) (Ambion, Thermo Fisher Scientific, Waltham, MA, US) were diluted in 100 µl of Opti-MEM medium (Gibco) plus 12 µl of HiPerFect reagent and incubated for 15 min at room temperature. Then, the mixture was added to the cells incubated in Opti-MEM Medium. The final siRNA concentration was 50 nM. After 3 h, 2 ml of DMEM supplemented with 0,1% FBS was added to the cells. 48 h later cells were stimulated with IGF-1 (10 ng/ml) for Ca^2+^ or proliferation assays.

A7r5 cells were transfected with miR-18a-5p mimic (Ambion, Thermo Fisher Scientific, Waltham, MA, US) and control miRNA mimic (Dharmacon, Lafayette, CO, US) using Lipofectamine^®^ RNAiMax Transfection Reagent (Thermo Fisher Scientific, Waltham, MA, US). Briefly, 5 µl lipofectamine was diluted in 250 µl Opti-MEM Medium, and miRNA mimics (Ambion, Thermo Fisher Scientific, Waltham, MA, US) were diluted in 250 µl of Opti-MEM Medium. The preparations were mixed in 1:1 proportion and incubated at room temperature for 5 min. After 3 h of incubation, 1 ml of DMEM supplemented with 5% FBS was added. The final miRNA mimic concentration was 30 nM. After transfection, cells were maintained in culture for 24 h for RNA assays or 48 h for protein assays and cytosolic Ca^2+^ measurements.

### Cell proliferation assay

Cell proliferation was estimated using the 5-bromo-2deoxyuridine (BrdU) incorporation assay. Coronary VSMC were seeded onto coverslips and grown to 70% confluence, cell growth was arrested by culturing in DMEM supplemented with 0.1% FBS and transfected with siRNA (siControl, siOrai1 or siSARAF). After 24 h, cells were incubated or not with IGF-1 (10ng/ml) for 48 h. BrDU (10µM) was added 18 h before immunostaining assays. Subsequently, cells were fixed with cold methanol for 5 min and incubated with 2M HCl for 30 min to denature the DNA. Cells were then washed twice with sodium tetraborate (0.1 M and pH 8.5) and blocked with a solution containing 3% heat-inactivated goat serum, 1% bovine serum albumin and 0.1% Triton X-100 in PBS1X. Next, cells were incubated with the primary antibody solution (anti-BrdU 1:300 dilution, Sigma-Aldrich, San Luis, MI, US) overnight at 4°C in a humidified chamber. After washing with PBS 1X, cells were incubated with the secondary antibody (1:400 AlexaFluor FITC 488 goat anti-mouse-IgG from Invitrogen, Waltham, MA, US) for 45 min in dark at room temperature followed by staining with DAPI (1:1000) for 5 min. The coverslips were finally mounted using Dako medium. Images were captured using an Olympus microscopy with the 20X objective equipped with an Olympus DP70 digital, camera and analyzed using *ImageJ s*oftware. All experiments were performed at least in triplicate, and for each experiment, 4-5 images were captured, and data were expressed as the percentage of BrdU-positive cells relative to the total number of cells (DAPI fluorescence).

### Dual RNA and Protein Isolation and Western Blotting of Protein fraction

Carotid arteries from the animal model were quickly frozen in liquid nitrogen and stored at −80°C. The arteries were lysed using TissuerLyser II (Qiagen, Hilden, Germany) with stainless steel beads. After homogenization, Qiazol Lysis reagent was added for RNA and protein extraction. Chloroform was added, then samples were vortexed and centrifuged. RNA was isolated from the aqueous phase using RNeasy Mini Kit (Qiagen, Hilden, Germany) as described below for arrays and qRT-PCR assays. Proteins were extracted from the organic and interphase fractions for western blotting. DNA was precipitated by adding 100% ethanol and centrifuging at 2000 g at 4°C. Proteins were precipitated using isopropanol and washed with guanidine hydrochloride (0.3 M in 95% ethanol). Each sample was resuspended in 50 µL of urea (10 M) and DTT (50 mM). The Bradford method was used for protein quantification.

For A7r5 cells, proteins were extracted using NP40 cell lysis buffer supplemented with a protease and phosphatase inhibitor cocktail (Roche, Basilea, Switzerland) and 1% PMSF incubated for 30 min on ice. Protein concentration was determined using a BCA Protein Assay Kit (Thermo Fisher Scientific, Waltham, MA, US). Equal amounts of protein (30-40 µg) were subjected to SDS-PAGE (10%-12,5% acrylamide) and electro-transferred to PVDF membranes. The membranes were blocked with 5% non-fat dry milk dissolved in Tris-buffered saline containing 0.1% Tween 20 (TTBS) for 2 h at room temperature. The membranes were then incubated overnight at 4°C with primary antibodies against mouse anti-Orai1 (1:250), mouse anti-STIM1 (1:1000), rabbit anti-SARAF (1:200), or mouse anti-αSMA (1:400) in TTBS containing 3% BSA. After washing, the membranes were incubated for 45 min at room temperature with horseradish peroxidase-anti-rabbit or anti-mouse conjugated anti-IgG (1:10000). Detection was performed using the Chemidoc Touch imaging system (Bio-Rad, Hercules, CA, US) with Clarity Western ECL or Clarity Max™ Western ECL reagents (Bio-Rad, Hercules, CA, US), and the images were analyzed using Image Lab Software.

### Co-immunoprecipitation assay

Carotid lysates were immunoprecipitated by incubation with 1 µg of anti-Orai1 binding to Protein A agarose beads (Merck Millipore, Germany) overnight at 4°C. The immunoprecipitates were resolved using 10% SDS-PAGE (Bio-Rad, Hercules, CA, US) and separated proteins were electrophoretically transferred onto nitrocellulose membranes and blocked for 1 h at room temperature. Blots were incubated overnight at 4°C with primary antibodies anti-SARAF antibody (1:1000) or anti-STIM1 (1:1000) in TTBS containing 3% BSA. To detect the primary antibody, the blots were incubated for 1 h with horseradish peroxidase-conjugated goat anti-rabbit IgG antibody at room temperature, and images were captured using a C-DiGit Chemiluminescent Western Blot Scanner (LI-COR Biosciences, Lincoln, NE, USA). After stripping, the blots were blocked overnight at 4°C and incubated with anti-Orai1 (1:1000) in TTBS with 3% BSA for 2 h at room temperature and then incubated for 1 h with horseradish peroxidase-conjugated goat anti-rabbit IgG antibody at room temperature.

### RNA isolation and qRT-PCR

Total RNA (mRNAs y miRNAs) was extracted using the RNeasy Mini Kit (Qiagen, Hilden, Germany) or the Nucleospin RNA II Kit (Macherey-Nagel, Duren, Germany), following the respective manufacturer protocols. RNA samples were quantified using NanoDrop™ while specific miRNAs quantification was performed using a Qubit 5.0 fluorometer (Thermo Fisher Scientific, Waltham, MA, US). Samples were stored at −80°C until their use. For mRNA analysis, 1 μg of total RNA was reverse transcribed into complementary DNA (cDNA) using iScript^TM^ Advanced cDNA Synthesis Kit (Bio-Rad, Hercules, CA, US), whereas miRNA reverse transcription was conducted using the miScript II RT kit (Qiagen, Hilden, Germany) according to the manufacturer’s instructions. qRT-PCR was performed in a FrameStar 384 Well PCR Plate (4titude, BIOKé, Leiden, the Netherlands). PCR mix includes SYBR Green reactive (iTaq™ Universal SYBR Green Supermix; Bio-Rad, Hercules, CA, US) and specific oligos for each gene or miRNA. For gene expression analysis, the following primers were used: Orai1 (sense: 5’-CCATAAGACGGACCGACAGT −3’, and antisense: 5’-GGGAAGGTGAGGACTTAGGC-3’), SARAF, sense 5’-GGACTCCTGTGGCTTGGTTA-3’ and antisense 5’-TGCTCTGTGGTCCTGTGAAG-3’; and 18S, (sense: 5’-AACGAGACTCTGGCATGCT-3’, and antisense: 5’-GCCACTTGTCCCTCTAAGA-3’). For miRNA expression, the following specific primers were used: Rn_miR-17a_1, Rn_miR-18a_1, Rn_miR-20a_1, Rn_miR-20b_1 (Qiagen, Germany). qRT-PCR was performed using an Applied Biosystems Viia7 7900HT thermocycler (Thermo Fisher Scientific, Waltham, MA, US) and data analysis was performed using Quant Studio RT PCR software. The expression levels of mRNA and miRNA were quantified using the threshold cycle (Ct) values, and relative expression was calculated using the comparative Ct method (2^−ΔΔCT^ method) normalized to 18S.

### Gene and miRNA transcriptomic analysis

The RNA was amplified and labeled using GeneChip®WT Pico Reagent Kit (Array (Thermo Fisher Scientific, Waltham, MA, US). RNA quality was assessed using an Agilent 2100 Bioanalyzer (Agilent Technologies, Santa Clara, CA, US), ensuring that the RNA Integrity Number (RIN) was greater than 7. The resulting cDNA was prepared for hybridization in the GeneChip® Clariom S Mouse Array (Thermo Fisher Scientific, Waltham, MA, US), while GeneChip^®^ miRNA 4.0 arrays (Thermo Fisher Scientific, Waltham, MA, US) was used to assess miRNA expressions. In both cases, the GeneChip® Fluidics Station 450 (Thermo Fisher Scientific, Waltham, MA, US) was used for washing and staining, and the GeneChip® Scanner 3000 was used for scanning. All procedures were carried out according to the manufacturer instructions. Data analysis was conducted using Transcriptome Analysis Console (TAC) v4.0 software (Thermo Fisher Scientific, Waltham, MA, US).

A comparative analysis was conducted between the different carotid groups and their respective controls. Genes with a fold-change greater than ±1.5, and miRNAs with a fold-change greater than ±2, and a p-value < 0.05 were considered differentially expressed.

Gene enrichment analysis was performed using g:profiler web tool (https://biit.cs.ut.ee/gprofiler/gost). Lists of genes significantly upregulated in 1 week compared to control or 1 week compared to 3 weeks were used as input. The resulting enriched Gene Ontology (GO) terms was processed and visualized as Dotplot using custom scripts in RStudio. For the heatmap representation, sample signal (Robust Multi-array Average, RMA) exported from TAC were used, and Z-score transformation was applied to each gene across samples.

The analysis of targeted genes predicted to be regulated by miRNAs was done using different bioinformatics resources: miRBD (miRDB v7.2, http://mirdb.org, Washington University Saint Louis, MO, US), TargetScan (Release 7.1, www.targetscan.org, Cambridge, MA, US) databases, and Exiqon tool application (http://www.exiqon.com/ microrna-target-prediction).

Functional enrichment analysis was performed using Gene Ontology (https://geneontology.org/) to identify pathways enriched in these genes.

### Luciferase Assay

Luciferase assay was performed using A7r5 smooth muscle cell line. The 2989 Kb sequence of the Orai1 gene promoter was synthesized and cloned in the pGL3-basic vector (Promega, Madison, WI, US) upstream of firefly luciferase by Condalab (Madrid, Spain). A7r5 cells were seeded in 24-well plates at 2.5×10^4^ cells/well in growth medium. 24 h later cells were transfected with 500 ng/well of pGL3-Orai1 luciferase constructs, 25 ng of the Renilla luciferase vector, pRL-TK (Promega, Madison, WI, US), and 30 nM of miR-18a-5p mimic, miR-20a-5p mimic, or control miRNA mimic using DharmaFECT kb transfection reagent (Dharmacon™, Lafayette, CO, US) according to the manufacturer instructions. After 24 h, cells were incubated with 5% FBS for another 24 h. Then, luciferase activity was examined using the Dual-Glo Luciferase Assay System and the luminescence was measured using a GloMax® Discover Microplate Reader (Promega, Madison, WI, US). Luciferase activities were expressed as relative Firefly/Renilla activities normalized to control. Four independent biological replicates were analysed.

### Statistical analysis

Data is presented as mean ± standard deviation (SD). Analysis of statistical significance was performed using GraphPad Prism v.8.4.3 (GraphPad Software, Inc. Group). Normal distributed variables were analyzed using *t* test Student for comparisons between two groups. The ordinary one-way ANOVA and multiple comparisons using *t* test without correction (Fisher’s LSD test) were performed when more groups were compared. For non-normally distributed variables we used the Mann-Whitney U test for comparisons between two groups and Kruskal-Wallis non-parametric test followed by Dunn’s post hoc test for multiple comparisons. Throughout the manuscript, *, **, ***, and **** indicate p values of <0.05, <0.01, <0.001, and <0.0001, respectively.

## RESULTS

### Balloon injury induces progressive neointima hyperplasia and impairs contractility

To characterize balloon-induced carotid injury, neointima hyperplasia was assessed at 1, 2 and 3 weeks after surgery, using classical hematoxiline-eosine staining. As shown in Figure 1A, progressive hyperplasia of the neointima was observed in the carotid artery sections following balloon injury compared to the control. Figure 1B and 1C confirm the progressive increase of the neointima/media ratio and the Carotid Intima-Media Area (CIMA), respectively, after balloon injury as compared to controls.

**Figure 1.**
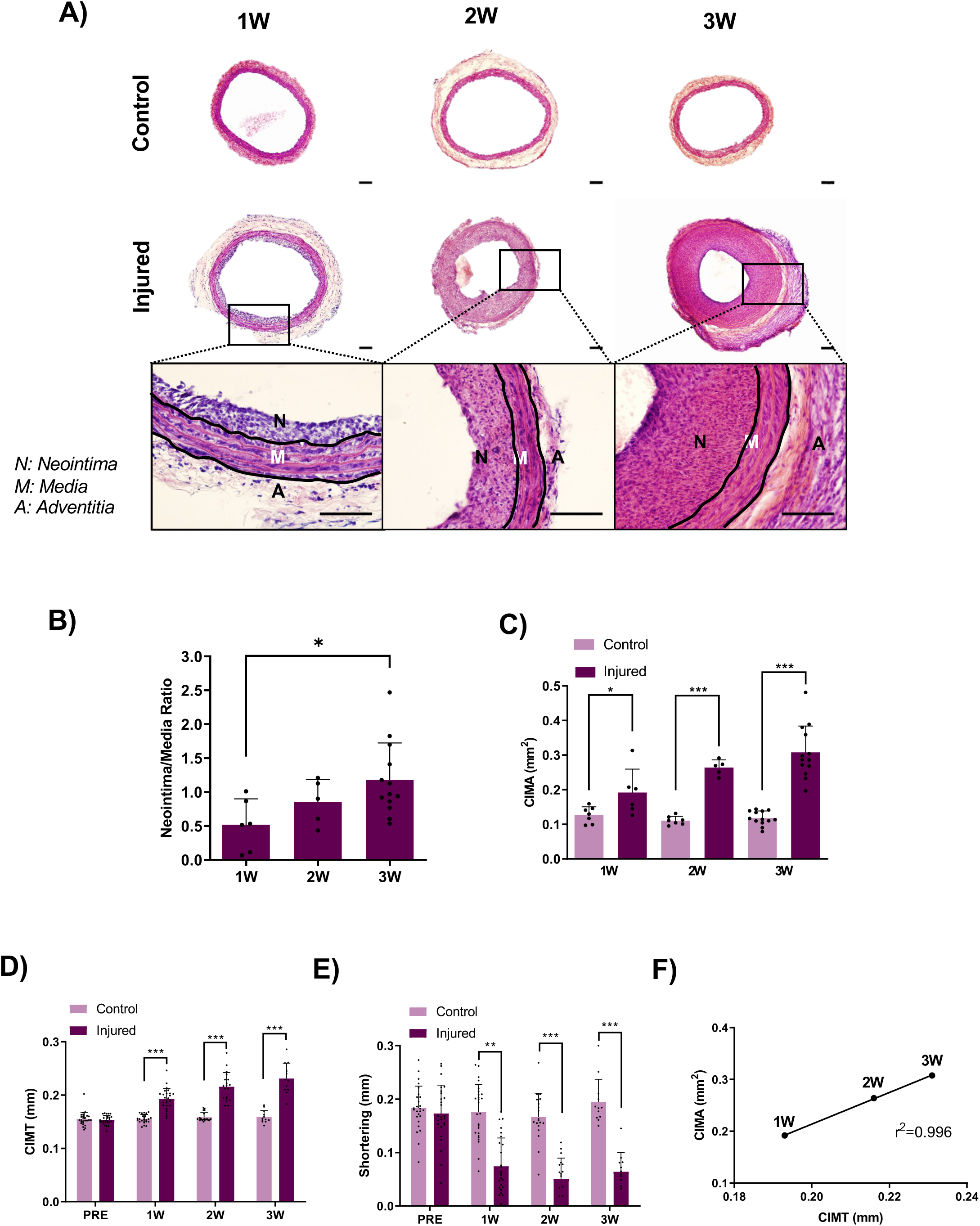
Neointima formation in rat carotid arteries after balloon injury. **A.** Representative images of hematoxilina-eosina staining of cross sections from control and injured carotid arteries (10X objective. Scale bar: 100 μm). **B.** Bar graph showing the Neointima/Media ratio of injured carotid arteries of hematoxilina-eosina staining of cross sections **C.** Bar graph indicates quantification of Carotid Intima-Media Area (CIMA) of injured carotid arteries compared with control. **D.** Bar graph showing Carotid Intima-Media Thickness (CIMT) by ultrasound before the intervention (PRE), in non-injured carotid artery (control) and in injured arteries 1 week, 2 and 3 weeks after angioplasty. **E.** Bar graph showing shortening performed by ultrasound under the same conditions than “D”. **F.** Correlation between CIMA measurements performed by histology and the CIMT evaluated by ultrasound in injured arteries. 1W: 1 week, 2W: 2 weeks, 3W: 3 weeks post-injury. Data are presented as mean ± SD. (*), (**), and (***) indicate significance with p < 0.05, p < 0.01, and p < 0.001, respectively, determined by one-way ANOVA. n = 3 independent experiments were conducted in each experiment.

To further validate balloon-induced carotid artery injury, endothelial denudation was assessed by immunofluorescence in cross-sections of control and injured carotid arteries. Supplementary Figure S1 shows a marked decrease in staining for the endothelial marker CD31 (red) in injured arteries compared with controls, persisting up to 3 weeks post-intervention, indicating substantial loss of the endothelial layer due to balloon-induced damage. αSMA staining (green) also confirms the formation of neointima by VSMC. Furthermore, we examined Carotid Intima-Media Thickness (CIMT) with ultrasound imaging, a method routinely used in clinical settings to predict the risk of coronary artery disease (*22*). Figure 1D shows that the CIMT was significantly higher in injured arteries compared to controls as early as the first week post-intervention. A significant decrease in arterial shortening was also observed at 1, 2, and 3 weeks post-intervention, indicating a significant reduction in the contractile capacity of injured arteries (Figure 1E; Supplemental video 1 and 2). Furthermore, CIMT, analyzed by ultrasound technique, and CIMA, evaluated in histological sections, shows a strong correlation (r^2^= 0.996), as indicated in Figure 1F.

Together, these findings demonstrate that balloon injury induces progressive neointimal hyperplasia in the carotid artery, driven by endothelial denudation and VSMC progression.

### Neointima formation is associated with significant transcriptomic dysregulation

To investigate the molecular basis of neointima formation following balloon injury, transcriptomic profiling was performed. Neointima formation is typically associated with complex alterations in gene expression, which contribute to vascular remodeling and repair. We performed a transcriptomic analysis on carotid arteries isolated from control and injured arteries at 1 and 3 weeks post-intervention. The analysis identified 879 and 449 differentially expressed genes (DEGs) between 1 week vs control, and 3 weeks vs control, respectively (Figure 2A-D; Fold change <1.5 or >1.5; *p* values < 0.05). Additionally, 1243 DEGs were detected between the 1 and 3 weeks injured samples, with 519 genes upregulated and 724 downregulated, indicating progressive changes in gene expression over time. Enrichment analysis of the DEGs revealed significant upregulation of biological processes and pathways involved in cell cycle regulation, migration, and proliferation, 1 week after the intervention, as illustrated in Figure 2E. The heatmap in Figure 2F confirms the upregulation of genes associated with proliferation 1 week post-intervention compared to control. In contrast, the expression of these genes decreased at 3 weeks after the intervention compared to 1 week timepoint (Figure 2G). Moreover, significant changes in the expression of Ca^2+^ associated channel, including those related to SOCE, was observed between 3 weeks and 1 week after injury (Figure 2H).

**Figure 2.**
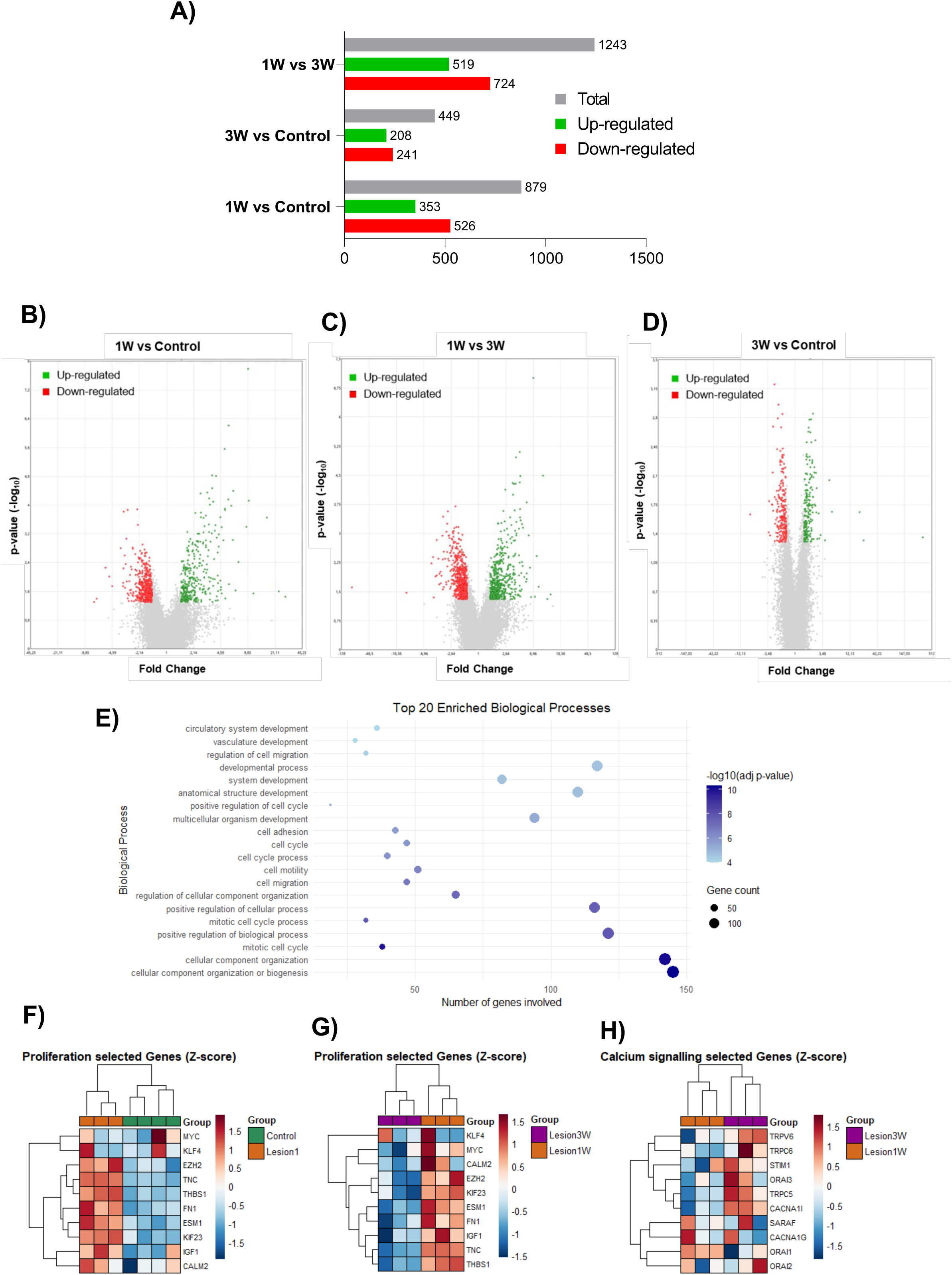
Neointima formation is associated with gene expression dysregulation. **A.** Representation of the total number of dysregulated genes (gray), downregulated genes (red), and upregulated genes (green) in injured arteries compared with controls. **B-D.** Volcano plots representation of the differentially expressed genes in injured carotid rats at 1 week compared with control **(B)**, 1 week compared with 3 weeks **(C)** and 3 weeks compared with control **(D)**. Upregulated genes are shown in green and downregulated genes are shown in red. Fold change: ± 1.5; p < 0.05. **E.** Dotplot showing the top 20 enrichment biological processes (Gene Ontology: Biological Process) identified using g:Profile, based on upregulated genes in the 1 week carotid injured after angioplasty compared with control carotids. The color represents the -log10 (adj. p-value) for each pathway, indicating the significance of the enrichment. Dots size reflects the number of genes (gene count) involved in each biological process. **F, G.** Heatmaps representing z-score expression values of proliferation related-genes at 1 week post-injury compared with control **(F)**, and 3 weeks post-injury compared with control **(G)**. **H.** Heatmaps representing z-score expression values of calcium signaling related-gene at 1 week (1W) in comparation with 3 weeks (3W) post-injury.

### Angioplasty-induced injury increases the expression of Orai1 and SARAF in the neointima

In line with the transcriptomic findings (Figure 2H) and with previous reports demonstrating the role of SOCE-dependent Ca^2+^ signaling in neointima formation, we analyzed the expression of Orai1 and SARAF in carotid artery following injury. Immunofluorescence analysis revealed that Orai1 and SARAF are expressed in the medial layer of control arteries (Figure 3A), whereas in injured arteries both proteins are predominantly expressed in the neointima layer up to 3 weeks after surgery. Additionally, a significant increase in the expression of Orai1 and SARAF are observed in injured arteries compared with controls, especially at 1 week, followed by a subsequent decline at 2 and 3 weeks post-intervention, as determined by western blotting (Figure 3B, 3C).

**Figure 3.**
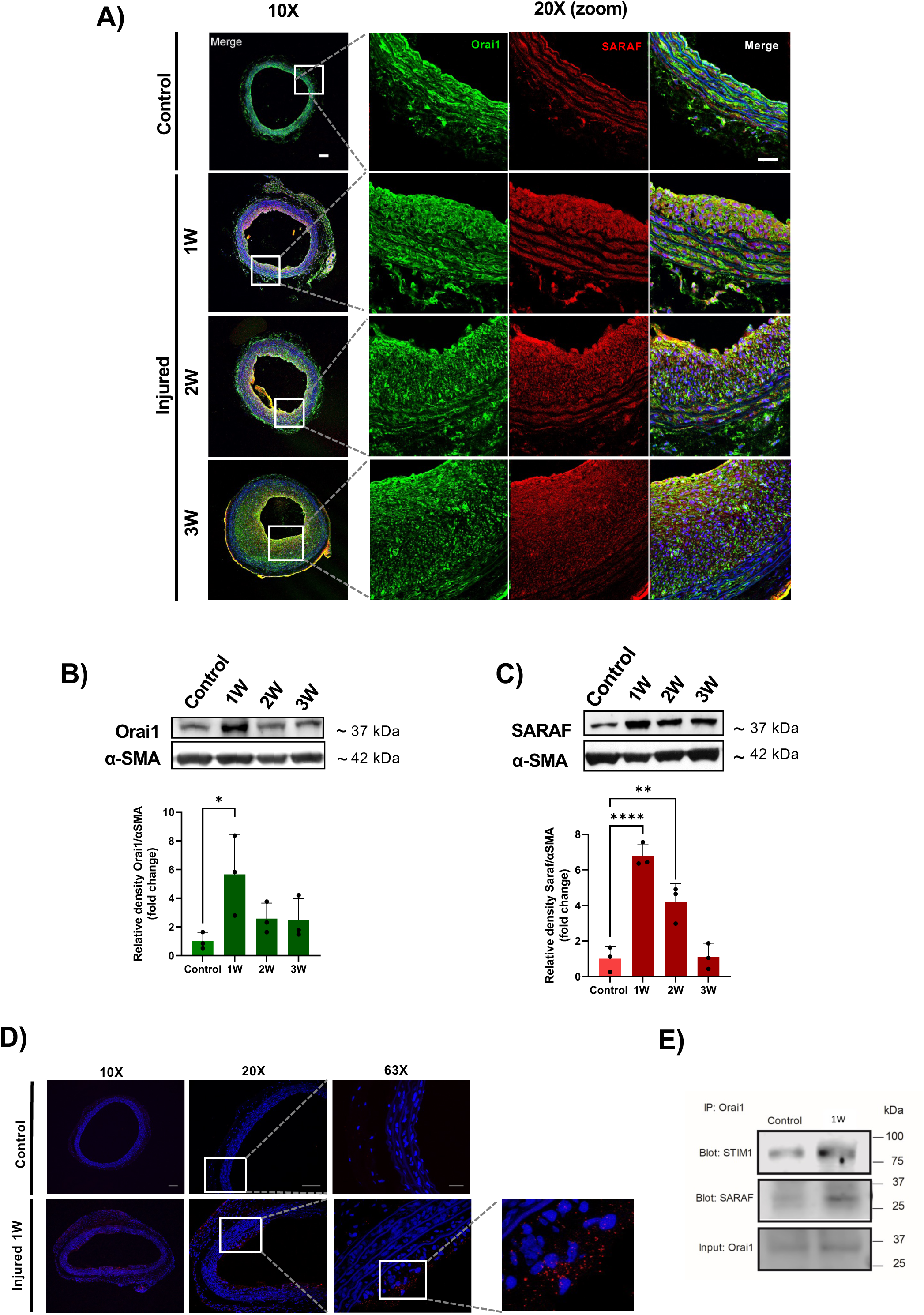
Orai1 and SARAF are upregulated in injured arteries. **A.** Representative images of immunofluorescence showing Orai1 (green), SARAF (red), and Dapi (blue) in a cross-section of the control and injured carotid arteries. Scale bar: 100 μm. **B, C.** Representative immunoblots (top) and summary data (bottom) showing the protein expression density of Orai1 **(B)**, and SARAF **(C)** normalized to its corresponding α-SMA. **D.** Representative images of Proximity ligation assay performed on a cross-section of control and injured arteries at 1 week post-surgery. Sections were incubated with mouse anti-Orai1 and rabbit anti-SARAF antibodies. Red fluorescent dots indicate proximity (< 40 nm) between Orai1 and SARAF. The nuclei were stained with DAPI (blue). Images were captured using a confocal microscope with 10X, 20X, and 63X objectives, with maximum z-projection of planes at 0.4 μm intervals. Scale bar: 100 μm. **E.** Co-immunoprecipitation of homogenized lysates from control and injured carotid arteries 1 week post-surgery with an anti-Orai1 antibody and analyzed by western blot using anti-STIM1 (Blot: STIM1), anti-SARAF (Blot: SARAF), and anti-Orai1 (Input: Orai1) antibodies as control. (n = pull of 5 arteries per conditions). 1W: 1 week; 2W: 2 weeks; 3W: 3 weeks post-injury. Data are presented as mean ± SD. (*), (**) and (****) indicate significance with p < 0.05, p < 0.01 and p < 0.0001, respectively.

Moreover, images in Figure 3A show prominent co-staining near the lumen of injured arteries, particularly at 2 and 3 weeks post-injury, suggesting colocalization of Orai1 and SARAF, as indicated by the yellow signal. To further investigate this, we analyzed the fluorescence patterns of Orai1 and SARAF as shown in Supplementary Figure S2. In control arteries, Orai1 staining exhibited showed a strong signal in the tunica media, while SARAF exhibited weaker signal. Interestingly, in injured arteries, both proteins displayed increased fluorescence intensity in the neointima layer near the lumen, where the fluorescence spectra of Orai1 and SARAF increasingly overlapped as the injury progressed. To further assess the possible interaction between Orai1 and SARAF, we used the PLA assay with Orai1 and SARAF antibodies. Figure 3D shows red puncta with highest density near the lumen in injured artery, indicating colocalization of both proteins in the neointimal region of arteries 1 week after injury. In contrast, no signal was detected in the control arteries. Furthermore, the co-immunoprecipitation assay shown in Figure 3E confirms that Orai1 interacts with SARAF in injured arteries. As expected, Orai1 also interacts with STIM1 in injured arteries. Notably, STIM1 protein is also upregulated in injured arteries 1 and 2 weeks after angioplasty compared with the control group (Supplementary Figure S3).

Altogether, these findings demonstrate the upregulation of SOCE-related proteins in injured arteries, suggesting a functional interaction between Orai1 and SARAF in the neointima.

### Orai1 and SARAF involvement in VSMC proliferation and SOCE

VSMC proliferation is crucial for neointima formation. Therefore, we examined proliferation in carotid artery sections using the proliferation marker Ki67. Figure 4A shows strong Ki67 staining in cells adjacent to the lumen, whereas staining significantly decreased in the rest of the neointima and media layers. Figure 4B confirms that the highest rate of proliferating cells occurs at 1 week post-surgery, followed by a prominent decrease over time. These results suggest that VSMC in the neointima near the lumen are more proliferative than those located deeper within the neointima or media.

**Figure 4.**
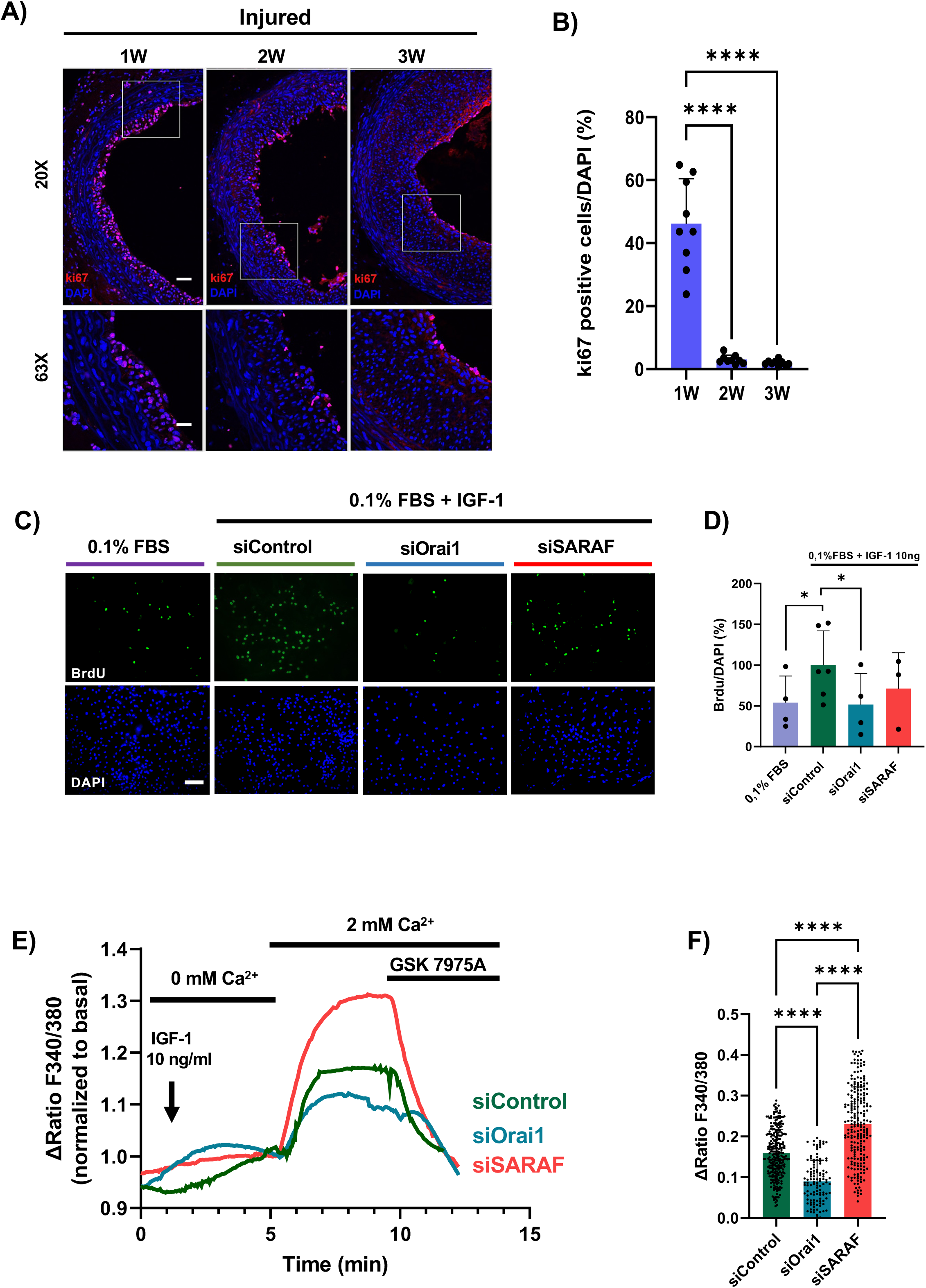
Orai1 and SARAF role in coronary artery VSMC proliferation and Ca^2+^ mobilization. **A.** Representative immunofluorescence images showing Ki67 (red) and DAPI (blue) staining in the cross-section of injured carotid arteries. Images were captured using a confocal microscope with 20X and 63X objectives. **B.** The bar graph represents the percentage of Ki67-positive cells relative to the total number of cells (DAPI) in the neointima. **C.** Representative immunofluorescence images taken with 20X objective showing BrdU (green) and DAPI (blue). Scale bar: 100 μm. **D.** Bar graph showing the percent of BrdU-positive cells relative to the total number of cells (DAPI) in coronary VSMCs in the presence of 0.1% FBS or 0.1% FBS with 10ng/ml of IGF-1. VSMC were transfected with siRNA control (siControl), siRNA of Orai1 (siOrai1), and siRNA of SARAF (siSaraf). **E, F.** Representative traces **(E)** and mean values **(F)** of 10 ng/ml IGF-1 induced Ca^2+^ influx in Fura-2 loaded coronary VSMCs. Recordings are from coronary VSMCs transfected with siRNA control (siControl), Orai1 siRNA (siOrai1), and SARAF siRNA (siSARAF). **F**. Δratio indicate the difference between the peak ratio after extracellular Ca^2+^ addition and its level immediately before the addition. GSK 7975A (GSK,10 μM) was applied to inhibit SOCE. (n=115-268 cells from 5 replicates). Data are expressed as mean ± SD. (*) and (****) indicate significance with p < 0.05and p < 0.0001, respectively.

Considering these findings, we explored the roles of Orai1 and SARAF in cells proliferation and Ca^2+^ mobilization using VSMC isolated from coronary arteries. Transcriptomic results in Figure 1J suggest that IGF-1 is upregulated 1 week after artery injury, thus we analyzed its effect on VSMC activation. Figures 4C and 4D show that IGF-1 (10 ng/ml) stimulation significantly increased the number of BrdU-positive cells, indicating enhanced VSMC proliferation. However, Orai1 siRNA attenuated IGF-1-induced proliferation, whereas SARAF siRNA had a minor effect. The efficacy of Orai1 and SARAF siRNAs in reducing their respective mRNA levels was confirmed as shown in Supplementary Figure 4.

To further explore the involvement of Orai1 and SARAF in IGF-1 responses, we examined their role in IGF-1-induced Ca²⁺ mobilization in VSMC. As shown in Figure 4E and 4F, the addition of 10 ng/ml IGF-1 in Ca^2+^-free solution evoked Ca^2+^ release and subsequently induced a significant increase in [Ca^2+^]_i_ after the re-addition of Ca^2+^. The induced Ca^2+^ influx was significantly decreased in cells transfected with Orai1 siRNA compared to controls, whereas SARAF siRNA potentiated IGF-1-evoked Ca^2+^ entry. The addition of SOCE inhibitor GSK 7975A (*23*) effectively decreased IGF-1-induced Ca^2+^ influx under all conditions, confirming the SOCE nature of the IGF-1-induced Ca^2+^ influx.

Taking together, these results confirm that Orai1 is essential for SOCE-mediated Ca^2+^ entry and contributes to coronary VSMC proliferation in response to IGF-1. In contrast, while SARAF does not appear to influence VSMC proliferation, it plays a modulatory role in Ca²⁺ mobilization.

### Neointima formation is associated with miRNA dysregulation

Given the importance of post-transcriptional regulation of gene expression during neointima hyperplasia, we examined the miRNA transcriptional profile in carotid artery at 1, 2, and 3 weeks post-intervention. As shown in Figure 5, the analysis of the hierarchical clustering (Figure 5A-5C) and volcano plot (Figure 5D-5F) indicate significant alterations in the expression of miRNAs (fold change ± 2 and p < 0.05). Specifically, we identified 1951, 1477, and 312 differentially expressed miRNAs in injured arteries compared with controls at 1, 2, and 3 weeks post-surgery, respectively (Figure 5G). Of these, 631, 441 and 88 miRNAs were upregulated at 1, 2, and 3 weeks post-surgery, respectively, whereas 1320, 1036, and 224 were downregulated at the corresponding time points. These findings demonstrate that the number of dysregulated miRNAs gradually decreases over time after arterial injury. Using the miRSystem tool and GO pathway enrichment analysis, we identified 745 biological pathways regulated by these genes. Among them, the most enriched pathways are notably involved in VSMC proliferation and migration, as summarized in Figure 5H.

**Figure 5.**
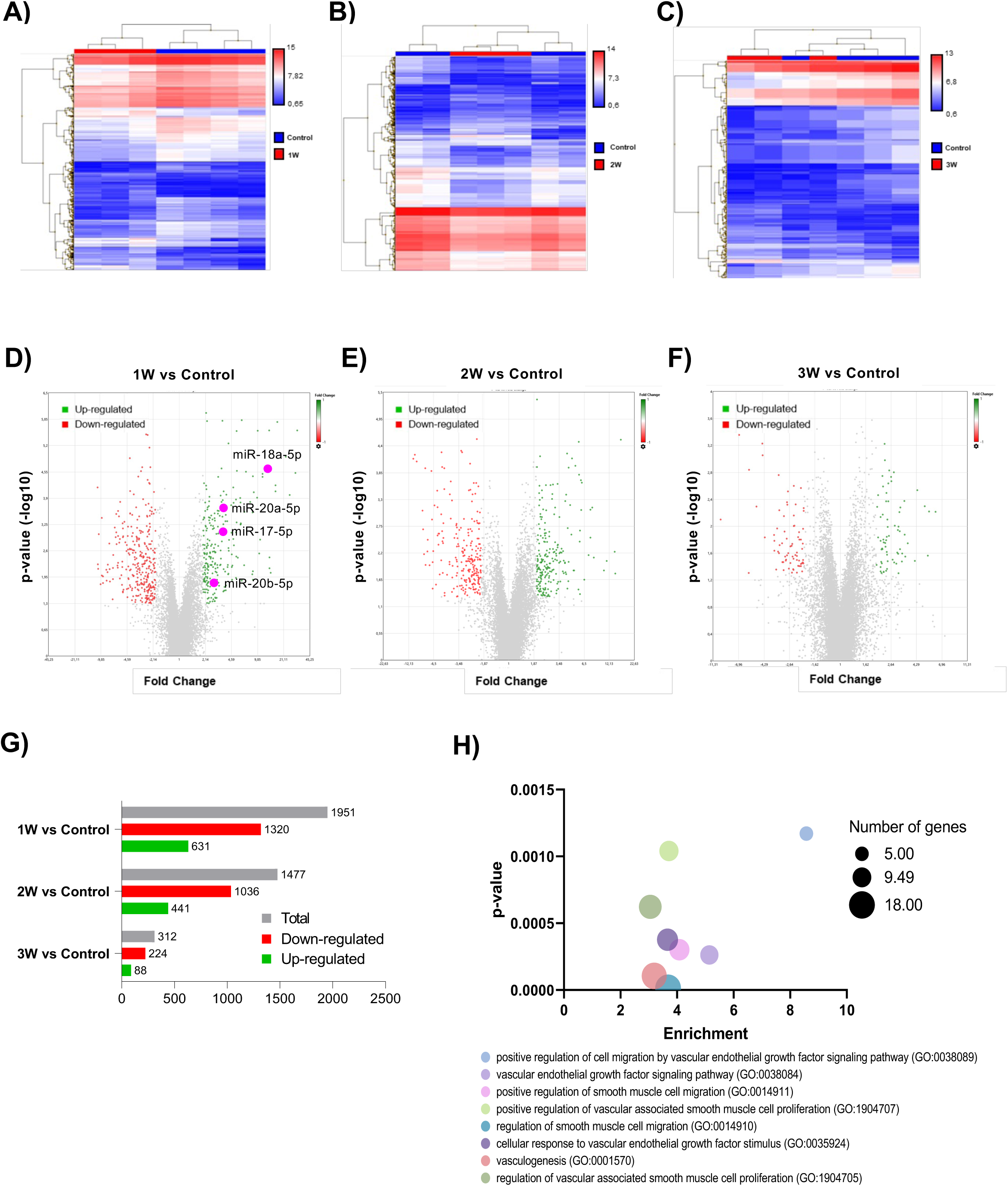
Microarray analysis of miRNA expression in injured and control carotid arteries. **A-C.** Heatmaps and representation of differentially expressed miRNAs profiles in injured carotid arteries (red) compared to control (blue) assessed at 1 week (1W), 2 weeks (2W), and 3 weeks (3W) post-injury. **D-F.** Volcano plots showing upregulated miRNAs in green and downregulated miRNAs in red in injured carotid rats compared with controls (Fold change = ± 2; p < 0.05). **G**. Representation of the total number of dysregulated miRNAs (gray), downregulated miRNAs (red), and upregulated miRNAs (green) in injured arteries compared with controls. **H**. Bubble plot enrichment representation of p-value for biological process pathways regulated by target genes of the miRNAs selected. The number of genes is indicated by the size of each bubble. Enrichment analysis identified 745 biological pathways.

### Orai1 is regulated by miR-18a-5p in VSMC

Given the dynamic changes in miRNA expression observed during vascular remodeling, we examined whether specific miRNAs contribute to the upregulation of Orai1 and SARAF. We performed an *in silico* analysis of miRNA array to identify those that may modulate their expression. In this context, several dysregulated miRNAs were identified as potential modulators of Orai1 and SARAF, as shown in Figures 6A and 6B. Among these miRNAs, we found a significant increase in the expression of miR-18a-5p (Fold change: 14.07), miR-20a-5p (Fold change: 3.73), miR-17-5p (Fold change: 3.53), and miR-20b-5p (Fold change: 2.72), all belonging to the miR-17-92 cluster, at 1 week post-surgery.

**Figure 6.**
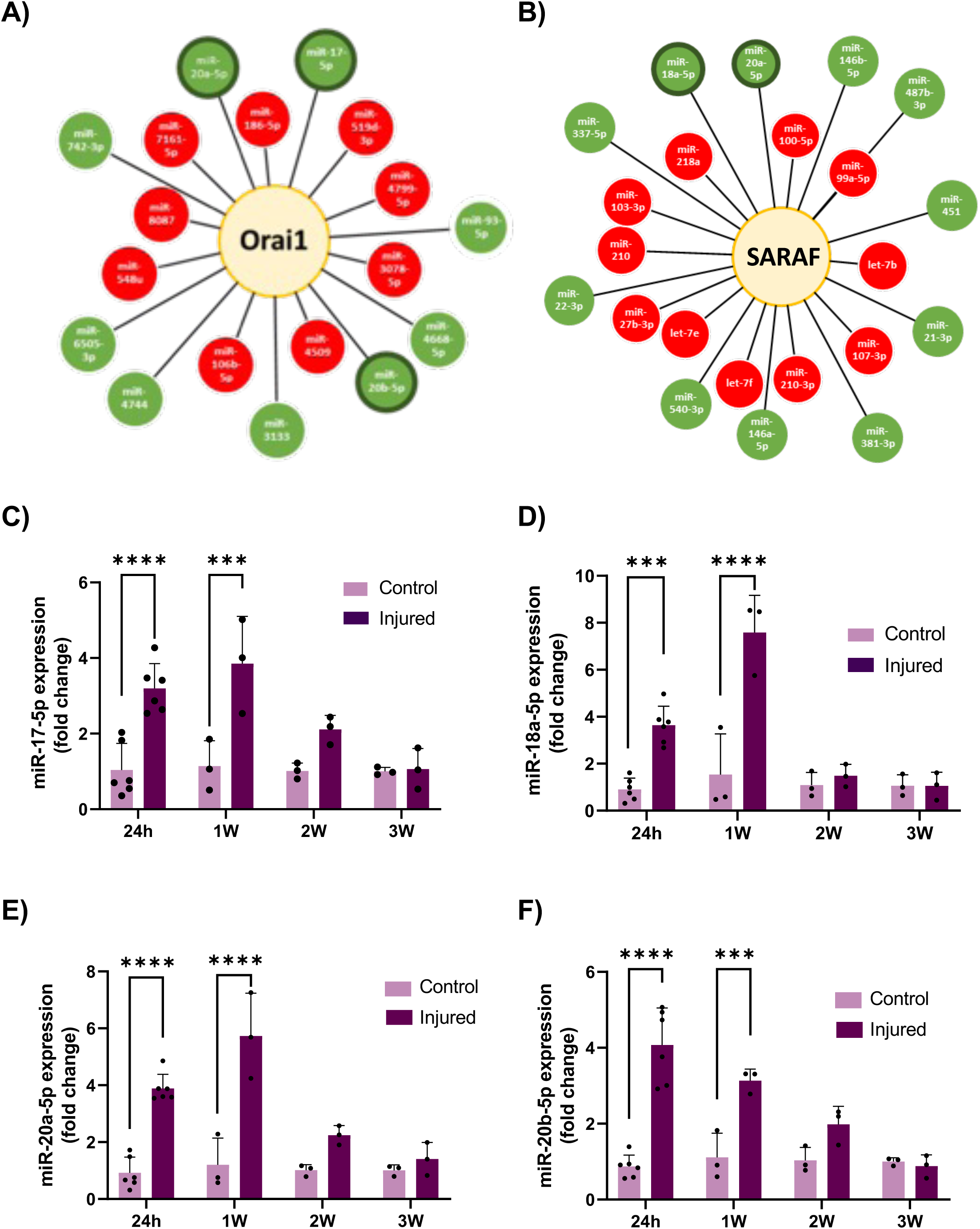
Expression of miRNA that target Orai1 in injured arteries. **A, B.** Diagram showing miRNAs differentially expressed in injured arteries compared with control at 1 week post-surgery that target Orai1 **(A)** and SARAF **(B)**. In red miRNAs with decreased expression in injured arteries. In green, upregulated miRNAs. Dark green outlines indicate miRNAs selected for their analysis in carotid arteries. **C-F.** Levels of miRNAs 17-5p **(C)**, 18a-5p **(D),** 20a-5p **(E)**, 20b-5p **(F)** in control and injured carotid arteries at 24 h, 1week (1W), 2 (2W), and 3 weeks (3W) after surgery, calculated using the 2^−ΔΔCt^ method normalized to the expression of the 18S. Values are presented as mean ± SD. (***), and (****) indicate significance with p < 0.001 and p < 0.0001, respectively.

We further examined the expression of these miRNAs by qRT-PCR in carotid arteries at different time points after injury. As shown in Figures 6C–6F, a significant increase in the expression of miR-17-5p, miR-18a-5p, miR-20a-5p and miR-20b-5p was observed at 24 hours and 1 week after injury compared to controls. Their expression levels gradually decreased at 2 and 3 weeks after arterial injury. Subsequently, we investigated whether miR-18a-5p, the most strongly upregulated miRNA from the 17-92 cluster in the injured arteries, could regulate the expression of Orai1 and SARAF. For these experiments, A7r5 VSMC were transfected with an miR18a-5p mimic, and mRNA levels of Orai1 and SARAF were analyzed. Figures 7A and 7B demonstrate that miR-18a-5p significantly increased *ORAI1* expression, while it tended to reduce *SARAF* expression.

**Figure 7.**
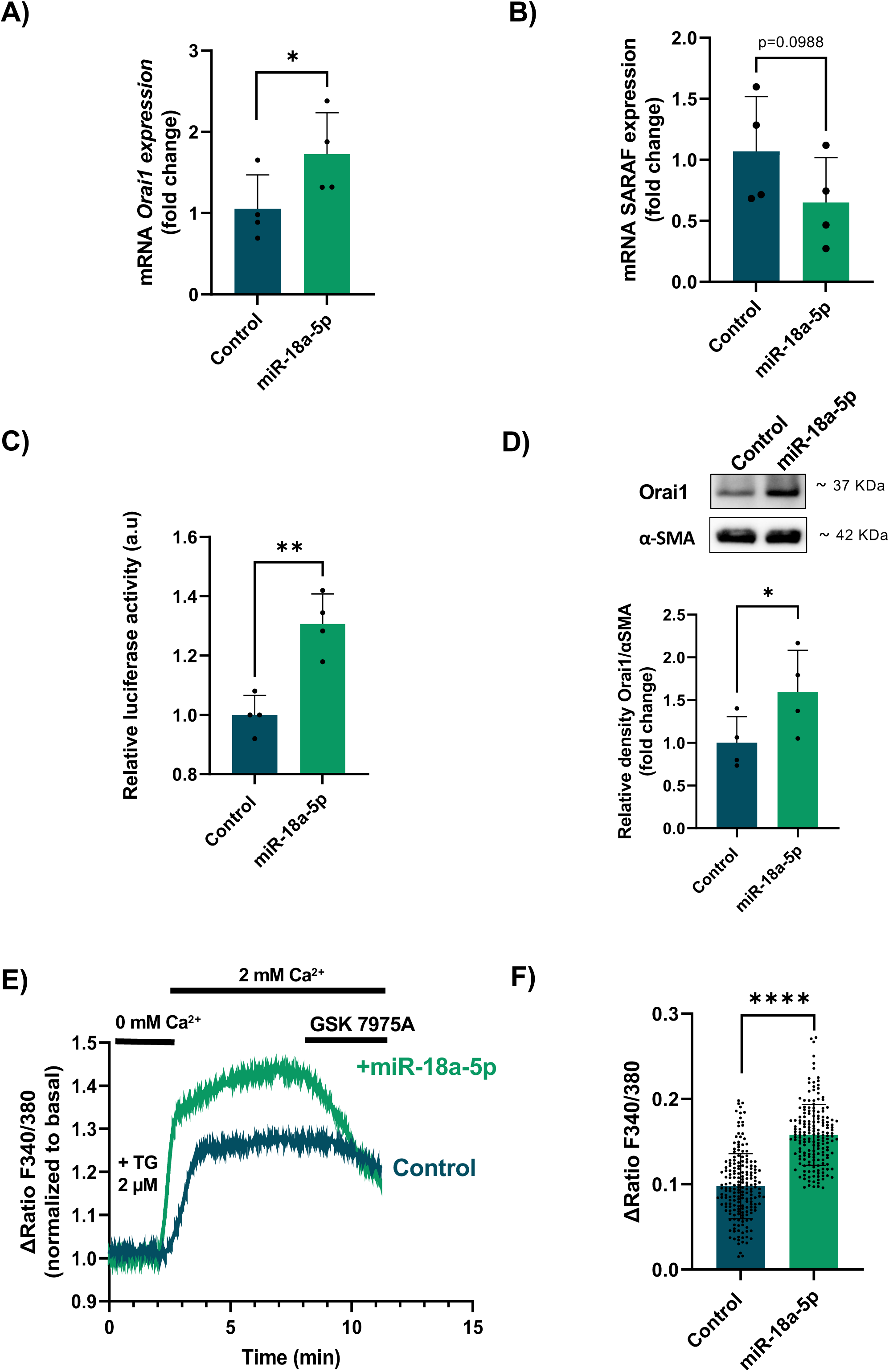
miR-18a-5p mimic stimulates Orai1 upregulation and potentiates SOCE in A7r5 cells. **A, B.** Bar graphs showing levels of Orai1 **(A)** and SARAF **(B)** mRNA in A7r5 cells transfected with mimic of miR18a-5p mimic (miR-18a-5p) and scrambled miRNA (Control). miRNA expression was calculated using the 2^−ΔΔCt^ method normalized to 18S expression. **C.** Relative Luciferase Assay upon transfection with pGL3 Basic 3Kb Orai1 and scrambled miRNA (Control), and pGL3 Basic 3Kb Orai1 and miR18a-5p mimic (miR-18a-5p) in A7r5 cells. **D.** Representative immunoblots (top) and summary data (bottom) showing the protein expression of Orai1 normalized to its corresponding α-SMA in A7r5 cells transfected with scrambled miRNA (Control) or miR18a-5p mimic (miR 18a-5p). Representative traces **(E)** and mean values **(F)** of thapsigargin-induced Ca^2+^ influx in Fura-2 loaded A7r5 cells. Recordings were acquired from A7r5 cells transfected with scrambled miRNA (Control) (n = 193), or miR-18a-5p mimic (miR-18a-5p) (n = 179), and preincubated with 2 µM of TG in Ca^2+^ free solution for 3 minutes. GSK 7975A (GSK,10 μM) was applied to inhibit SOCE. **F.** Δratio indicate the difference between the peak ratio after extracellular Ca^2+^ addition and its level immediately before the addition. (n = 4 cell culture). Values are presented as mean ± SD. (*), (**), and (****) indicate significance with p < 0.05, p < 0.01, and p < 0.0001, respectively.

Considering these findings, along with our earlier results showing the upregulation of both Orai1 and miR-18a-5p in injured arteries, we investigated whether miR-18a-5p directly stimulates *ORAI1* transcriptional activity, potentially leading to enhanced SOCE. Figure 7C shows that A7r5 cells co-transfected with the miR-18a-5p mimic and a reporter plasmid containing the rat *ORAI1* promoter driving firefly luciferase activity exhibited significantly increased luciferase activity, indicating enhanced *ORAI1* transcriptional activation. Moreover, as shown in Figure 7D, miR-18a-5p further increased Orai1 protein expression.

To determine whether miR-18a-5p-mediated Orai1 upregulation enhances SOCE, we examined the effect of thapsigargin on Ca^2+^ influx in cells transfected with miR-18a-5p mimic. Figures 7E and 7F show that in A7r5 cells incubated with thapsigargin, the re-addition of extracellular Ca^2+^ led to significantly greater Ca^2+^ influx in cells transfected with miR-18a-5p compared to control. The evoked Ca^2+^ influx was inhibited by GSK 7975A, confirming that it was mediated by SOCE. The analysis of cell responses revealed that 179 of 265 (68%) miR-18a-5p-transfected cells displayed a Ca^2+^ increase with a mean amplitude that was substantially higher than that observed in control cells, as summarized in Figure 7F. These results indicate that a greater proportion of miR-18a-5p-transfected cells exhibit exacerbated SOCE. Thus, these data suggest that miR-18a-5p, which is highly expressed in injured arteries, stimulates Orai1 upregulation and consequently enhances Ca²⁺ influx.

Finally, to extend the main findings to human arteries, we analyzed Orai1expression in human coronary arteries obtained from explanted ischemic hearts and in carotid arteries from healthy donors. As shown in Figure 8A, mRNA levels are significantly elevated in the ischemic arteries compared to healthy controls. Moreover, Figure 8B shows positive Orai1 staining in the medial layer in the ischemic arteries.

**Figure 8.**
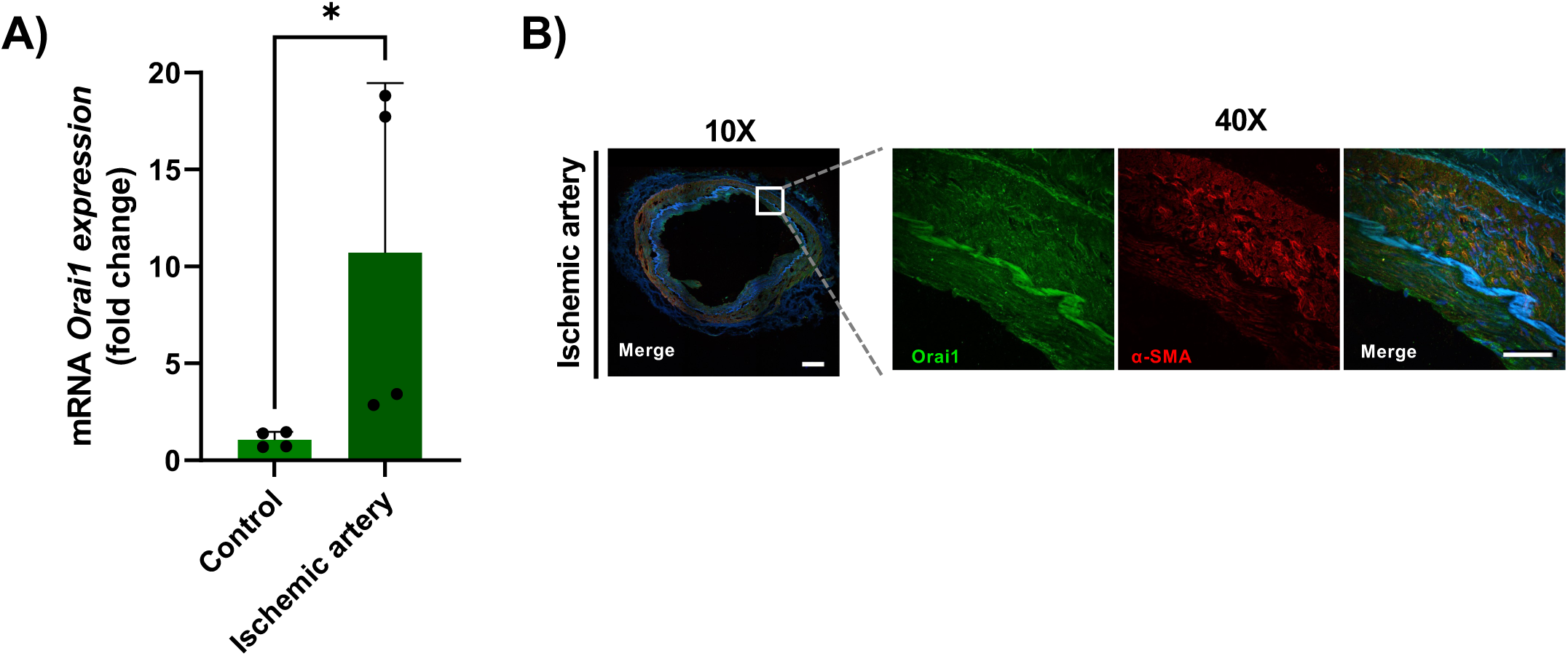
Expression of Orai1 in human arteries. **A.** mRNA levels of Orai1 in human coronary arteries obtained from explanted ischemic hearts and carotid arteries from healthy donors using the 2^−ΔΔCt^ method normalized to 18S expression. **B**. Representative images of immunofluorescence showing Orai1 (green), α-SMA (red) and Dapi (blue) in a cross-section of human coronary arteries from ischemic hearts. Images were captured using a confocal microscope with 10X and 40X objectives. Scale bar: 100 μm.

## DISCUSSION

Restenosis following successful vessel angioplasty remains a major clinical challenge, despite ongoing advances in medical technologies. In this study, we employed the well-established rat carotid angioplasty model to investigate the molecular mechanisms underlying neointima formation after vascular injury. Our findings confirm the critical role of SOCE-related proteins in VSMC proliferation and subsequent neointimal development, such as Orai1 and SARAF. In addition, we identified several dysregulated miRNAs from the miR-17-92 cluster following injury, which target SOCE-related proteins. Among them, miR-18a-5p was found to stimulate Orai1 upregulation, thereby enhancing SOCE activity. These findings identify a novel miR-18a-5p/Orai1 regulatory axis that contributes to VSMC proliferation and neointima formation following vascular injury.

Using ultrasound imaging to assess Carotid Intima-Media Thickness (CIMT) enabled the early detection of progressive neointima formation prior to histological analysis. Our analysis revealed a strong and reliable positive correlation between CIMT measurements obtained by ultrasound and Carotid Intima-Media Area (CIMA) values derived from histological analysis. These findings validate ultrasound imaging as a reliable, non-invasive method for monitoring vascular remodeling. This approach offers a significant advantage in preclinical research by reducing the number of experimental animals required and facilitating the accurate determination of optimal time points for their tissue collection.

Neointima formation is driven by complex gene expression alterations that contribute to vascular remodeling. Transcriptomic analysis of differentially expressed genes revealed significant dysregulation of biological processes and signaling pathways associated with cell cycle regulation and proliferation following vascular injury, among other relevant cellular pathways. Notably, genes involved in Ca^2+^ homeostasis, including those related to SOCE, exhibited marked expression changes between 1 and 3 weeks post-injury. Heatmap analysis indicated upregulation of Orai1 at 1 week compared to 3 weeks, and protein-level analysis revealed a significant increase in Orai1, STIM1 and SARAF expression as early as 1 and/or 2 weeks post-injury. In fact, in agreement with previous reports (*24*, *25*), we observed significant upregulation of Orai1 and STIM1 in the neointima and carotid artery sections.

We also observed increased SARAF expression, whose role in neointima formation remains unclear, within the same timeframe. Interestingly, Orai1 and SARAF signals were predominantly co-localized in the neointima, and their interaction was confirmed by co-immunoprecipitation and PLA, suggesting that SARAF may modulate Orai1 activity during the early stages of neointima formation. Strong staining of Orai1 and SARAF near the lumen, a region with high proliferative activity as indicated by Ki67 staining, further supports their involvement in VSMC proliferation. In addition, consistent with its classical inhibitory effect on SOCE (*6*), SARAF silencing led to significant SOCE activation. We therefore expected that SARAF might also stimulate VSMC proliferation. However, while Orai1 silencing significantly reduced IGF-1-induced coronary VSMC proliferation, SARAF silencing had no significant effect, suggesting a primary role for Orai1 in this context. This finding contrasts with recent studies demonstrating that SARAF can interact with Orai1 to facilitate SOCE activation and promote proliferation in other cell line (*21*, *26*), supporting a dual regulatory role for SARAF in both activation and inactivation of Orai1 (*7*). Recently, SARAF overexpression has been associated with decreased neointimal hyperplasia through suppression of VSMC proliferation and migration, via inhibition of STIM1 and Orai1 expression at 4 weeks post-injury (*8*). These discrepancies highlight the need for further investigation to clarify the precise role of SARAF in VSMC activation and neointimal hyperplasia. Importantly, herein we showed for the first time that Orai1 is overexpressed in human coronary arteries isolated from ischemic heart tissue compared with healthy human carotid artery, suggesting its clinical relevance.

Although Orai1 upregulation following vascular injury has been demonstrated previously, the precise molecular mechanisms driving its increased expression remain poorly understood. Recent studies have focused on the role of epigenetic regulators, particularly miRNAs, in modulating gene expression in cardiovascular diseases. This study revealed significant dysregulation of miRNAs at 1, 2, and 3 weeks post-injury, with the highest number of altered miRNAs detected at 1 week post-injury, likely reflecting an early inflammatory response triggered by vascular intervention (*27*). Further analysis confirmed that these miRNAs participate in cell cycle progression and proliferation, hallmarks of restenosis and neointima development. Moreover, *in silico* analysis identified Orai1 and SARAF as potential targets of miRNAs from the miR-17-92 cluster, a well-known implicated in both cancer and cardiovascular diseases (*28*, *29*). Herein, we observed sustained overexpression of miR-17-5p, miR-18a-5p, miR-20a-5p, and miR-20b-5p in injured arteries from 24 hours to 1 week post-surgery, coinciding with peak VSMC proliferation and the upregulation of Orai1 and SARAF. Our study reveals the unusual finding that miR-18a-5p enhances Orai1 promoter activity and expression. Although miRNAs typically repress gene expression, instances of indirect upregulation have been reported (*16*, *30*). To date, only a few miRNAs, such as miR-519 and miR-93-5p, have been reported to negatively regulate Orai1 expression (*12*, *31*). Our results provide the first evidence that miR-18a-5p, whose expression is markedly increased after injury, positively regulates Orai1 expression. Specifically, a luciferase reporter assay demonstrated that miR-18a-5p activated the Orai1 promoter, leading to increased levels of Orai1 protein. This upregulation was accompanied by enhanced thapsigargin-induced SOCE in VSMC, confirming the functional relevance of Orai1 as a miR-18a-5p target. Of note, miR-18a-5p barely affected SARAF expression, suggesting certain degree of target specificity.

The precise mechanism by which miR-18a-5p mediates this effect in VSMCs warrants future investigation. Signaling pathways such as AKT/ERK, known to be activated by miR-18a-5p in other contexts (*32*) and can regulate gene transcription, represent plausible candidates for subsequent mechanistic studies. Recently, we demonstrated that CREB activation enhances Orai1 promoter activity and expression in myocardial cells (*17*), supporting a potential indirect mechanism.

Altogether, the current study reveals a previously uncharacterized mechanism during neointima formation, involving miR-18a-5p modulation of Orai1. Identifying such regulatory miRNAs and elucidating their networks could provide novel insights into the epigenetic control of SOCE-related protein and may offer new therapeutic avenues for restenosis following vessel injury.

## ACKNOWLEDGMENTS

This research is part of the projects PID2022-136279NB-C21 and PID2022-136279NBC22 funded by MICIU/AEI/10.13039/501100011033/ and by “ERDF a way of making Europe”, and Junta de Andalucia-[proyexcel_00530; US-1381135]. M M-B and I G-O received a fellowship from “Fondation Bergonié”. D F-B is supported by EMERGIA talent followship. B D-L was supported by a contract from AEI/Spanish Ministry of Science and Innovation and INVESTIGO, respectively. The authors wish to thank F.J. Moron for advice and technical assistances in experiments at IBiS core facilities.

---

**Supplementary figure S1.**
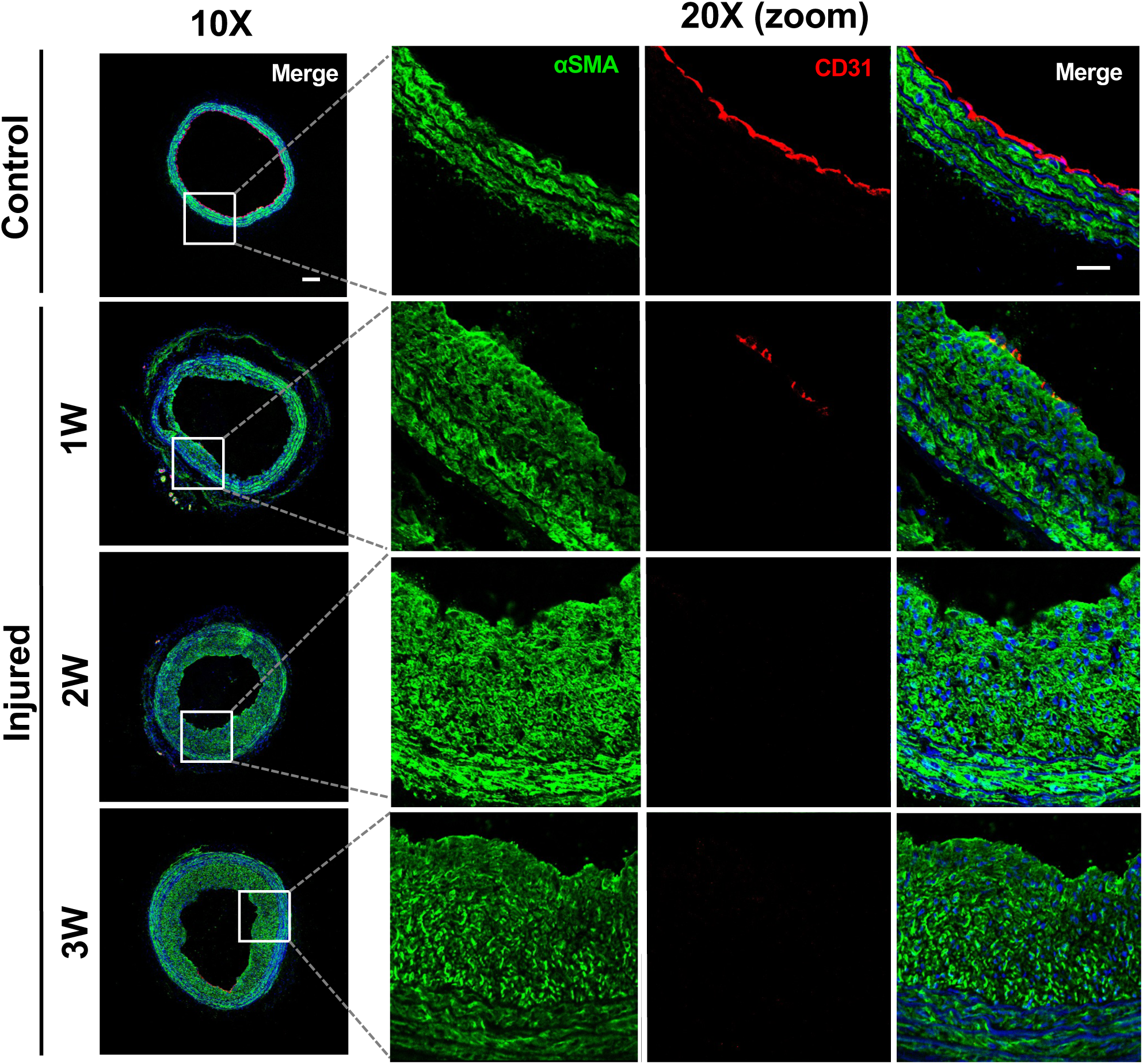
Endothelial denudation of injured carotid arteries. Representative images of immunofluorescence staining of cross sections from control and injured carotid arteries at 1, 2 and 3 weeks post-injury showing smooth muscle cells in intima and neointima labeled with αSMA (green) and endothelial cells with CD31 (red). Images were captured using a confocal microscope with 10X and 20X objectives. The magnification is 2X with the 20X objective. 1W: 1 week; 2W: 2 weeks; 3W: 3 weeks post-injuy. Scale bar, 100 μm.

**Supplementary figure S2.**
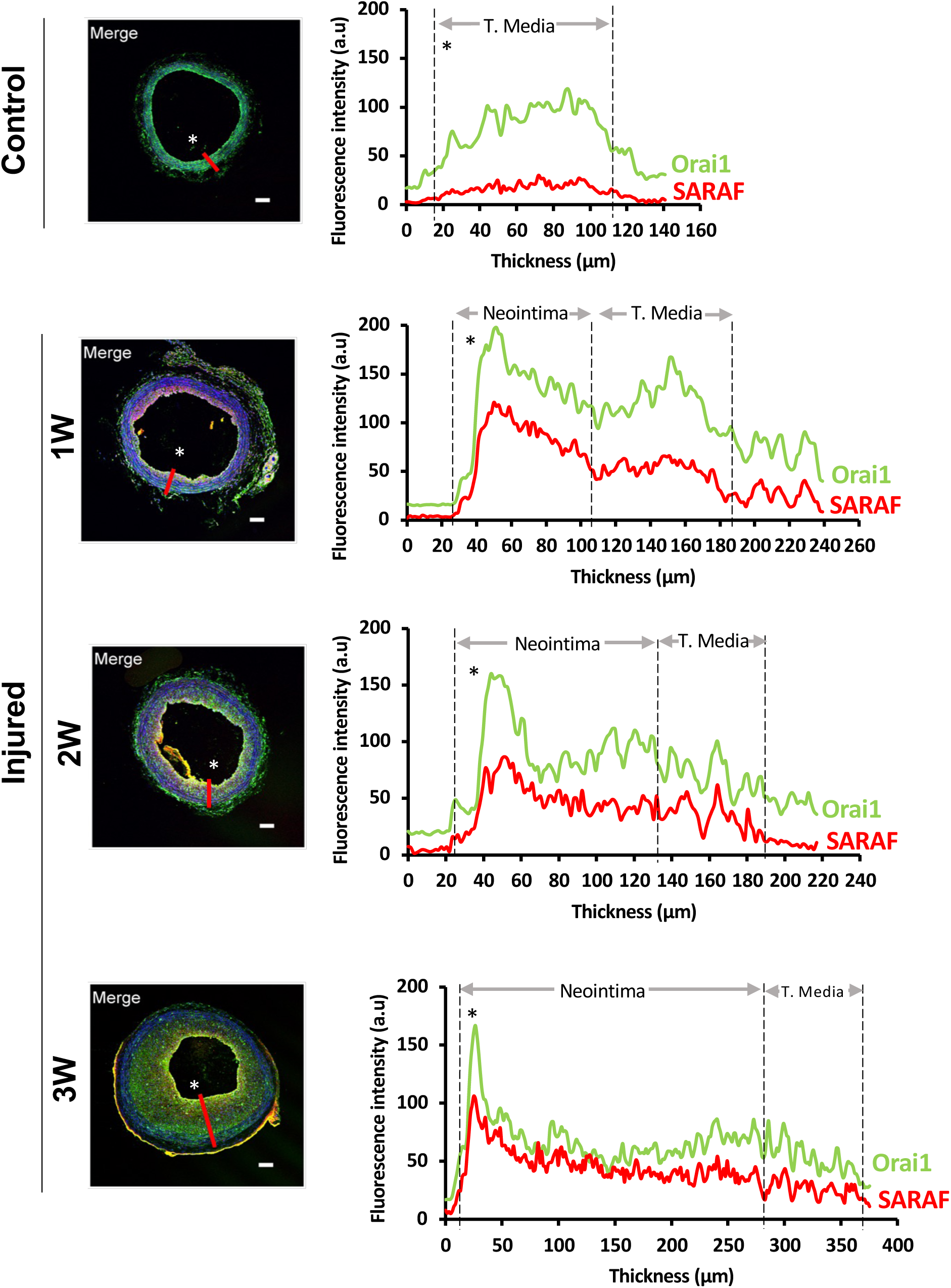
Fluorescence intensity of Orai1 and SARAF in control and injured carotid arteries following angioplasty. Representative images of immunofluorescence of control and injured carotid arteries at 1, 2 and 3 weeks post-surgery showing Orai1 (green), SARAF (red) and Dapi (blue). Images were captured using a confocal microscope with 10X objective. In right, graphs representing Orai1 and SARAF fluorescence intensity profiles across a line drawn through the thickness of the tissue was compared in the media and neointima layers. *Indicate the region near the lumen. 1W: 1 week; 2W: 2 weeks; 3W: 3 weeks post-injuy. Scale bar, 100 μm.

**Supplementary figure S3.**
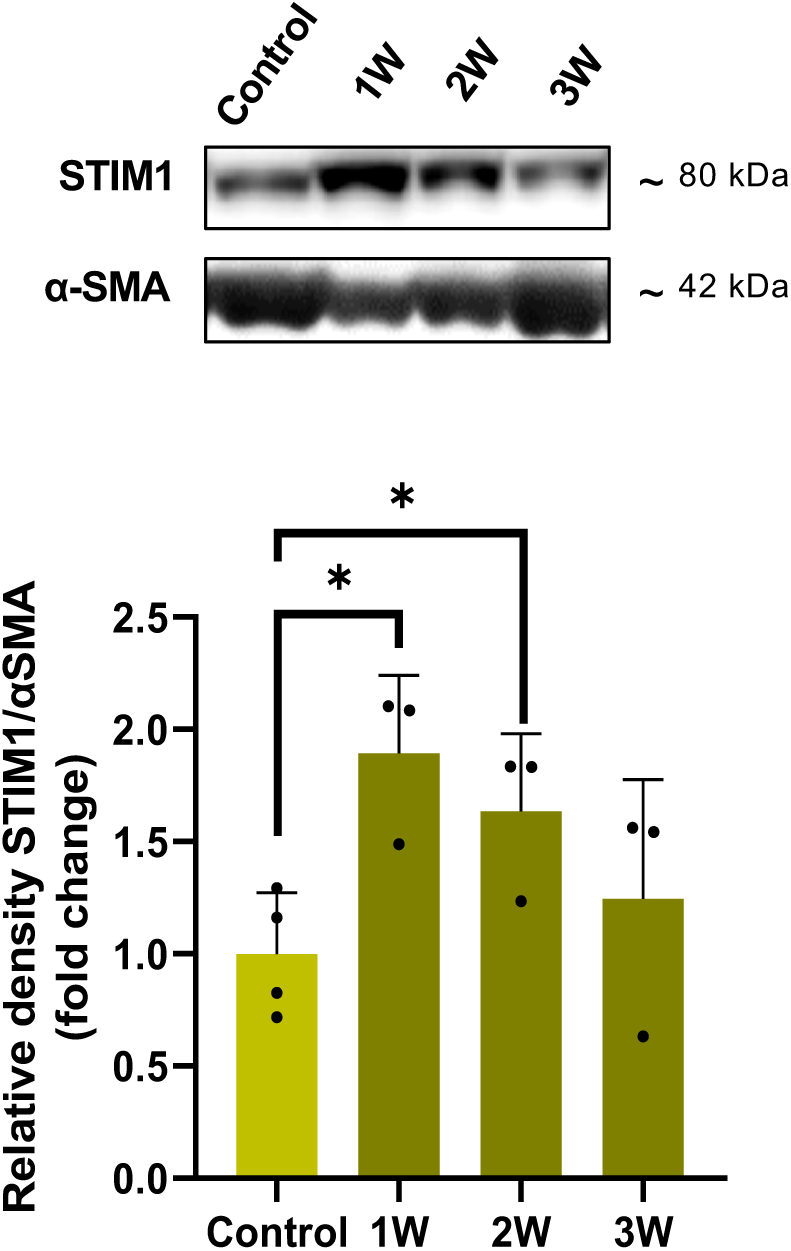
STIM1 is upregulated in injured arteries. Representative immunoblots (top) and summary data (bottom) showing the protein expression density of STIM1 normalized to its corresponding α-SMA in control and injured arteries. Data are presented as mean ± SD. (*) indicates significance with p < 0.05.

**Supplementary figure S4.**
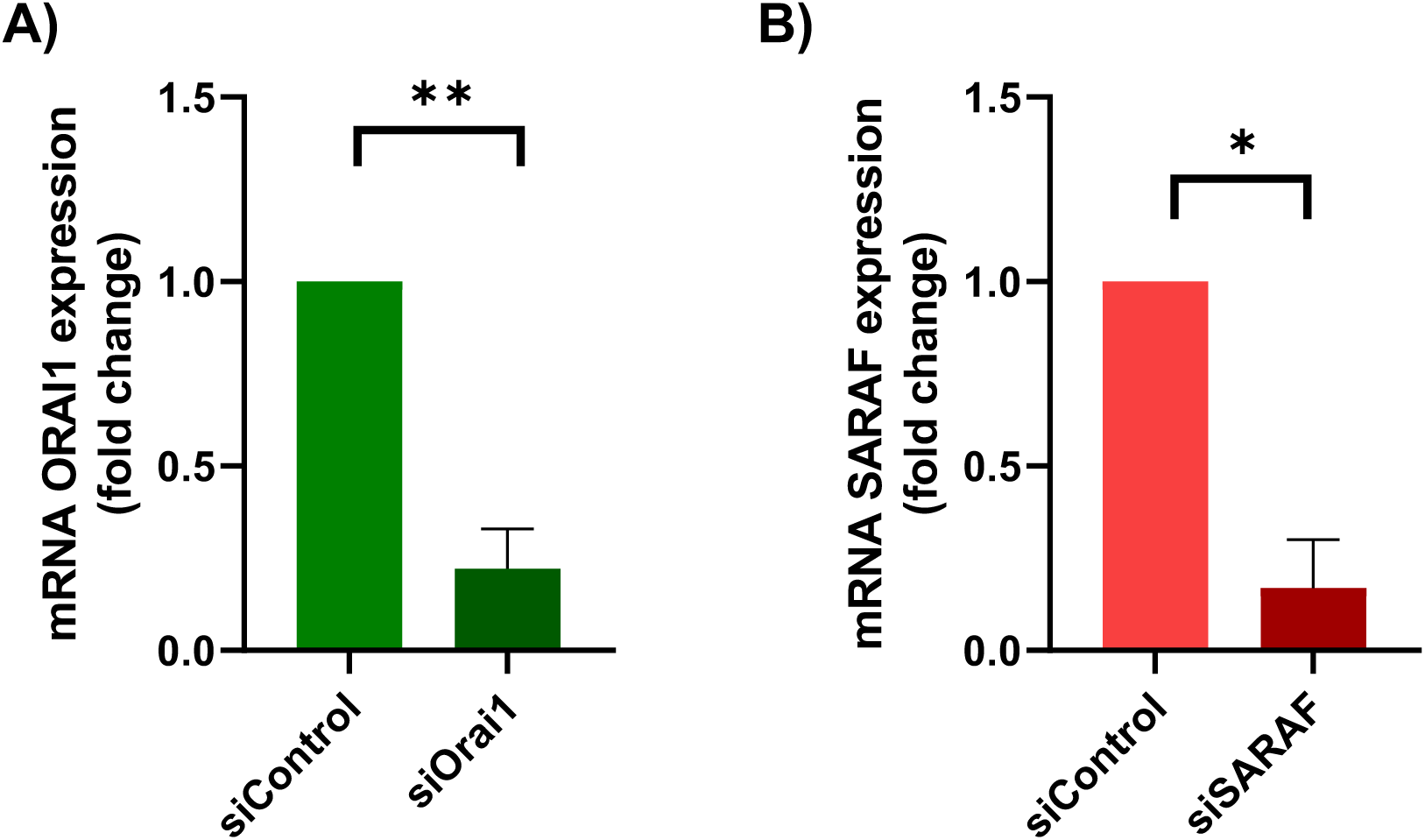
Silencing of Orai1 and SARAF in primary culture of coronary smooth muscle cells (VSMCs). mRNA expression levels of Orai1 **(A)** and SARAF **(B)** measured by RT-qPCR in cells transfected with the siRNA of Orai1 (siOrai1) or siRNA of SARAF (siSARAF) compared with those transfected with the scrambled RNA (siControl) (n = 3). Values were calculated using the 2^-ΔΔCt^ normalized to the expression of 18S. Values are presented as mean ± SD. (*) and (**) indicate significance with p < 0.05 and p < 0.01, respectively.

